# Whole Genome Comparison of Pakistani Corona Virus with Chinese and US Strains along with its Predictive Severity of COVID-19

**DOI:** 10.1101/2020.05.01.072942

**Authors:** Rashid Saif, Tania Mahmood, Aniqa Ejaz, Saeeda Zia, Abdul Rasheed Qureshi

**Affiliations:** Institute of Biotechnology, Gulab Devi Educational Complex, Lahore, Pakistan; Decode Genomics, 323-D, Punjab University Employees Housing Scheme (II), Lahore, Pakistan; Department of Sciences and Humanities, National University of Computer and Emerging Sciences, Lahore, Pakistan; Out-Patient Department-Pulmonology, Gulab Devi Chest Hospital, Ferozepur Road, Lahore, Pakistan

**Keywords:** Pakistani SARS-nCoV2, Phylogenetic analysis, Variant calling pipeline, 3D structural modelling

## Abstract

Recently submitted 784 SARS-nCoV2 whole genome sequences from NCBI Virus database were taken for constructing phylogenetic tree to look into their similarities. Pakistani strain MT240479 (Gilgit1-Pak) was found in close proximity to MT184913 (CruiseA-USA), while the second Pakistani strain MT262993 (Manga-Pak) was neighboring to MT039887 (WI-USA) strain in the constructed cladogram in this article. Afterward, four whole genome SARS-nCoV2 strain sequences were taken for variant calling analysis, those who appeared nearest relative in the earlier cladogram constructed a week time ago. Among those two Pakistani strains each of 29,836 bases were compared against MT263429 from (WI-USA) of 29,889 bases and MT259229 (Wuhan-China) of 29,864 bases. We identified 31 variants in both Pakistani strains, (Manga-Pak vs USA=2del+7SNPs, Manga-Pak vs Chinese=2del+2SNPs, Gilgit1-Pak vs USA=10SNPs, Gilgit1-Pak vs Chinese=8SNPs), which caused alteration in *ORF1ab, ORF1a* and *N* genes with having functions of viral replication and translation, host innate immunity and viral capsid formation respectively. These novel variants are assumed to be liable for low mortality rate in Pakistan with 385 as compared to USA with 63,871 and China with 4,633 deaths by May 01, 2020. However functional effects of these variants need further confirmatory studies. Moreover, mutated N & ORF1a proteins in Pakistani strains were also analyzed by 3D structure modelling, which give another dimension of comparing these alterations at amino acid level. In a nutshell, these novel variants are assumed to be linked with reduced mortality of COVID-19 in Pakistan along with other influencing factors, these novel variants would also be useful to understand the virulence of this virus and to develop indigenous vaccines and therapeutics.

## Introduction

The SARS pandemic engendered new avenues to ponder and identify variations in this animal based virus Severe Acute Respiratory Syndrome novel Corona virus 2 (SARS-nCoV2) that how human receptor Angiotensin-converting enzyme 2 (ACE2) become ideally compatible with the spike region of this virus and as a result COVID-19 spread human population globally [1]. In the current century, the first wave of transmission started from SARS CoV in Guangdong, China and thereafter disseminated worldwide which resulted in 916 fatalities [2]. Next in 2012, Middle East Respiratory Syndrome Corona virus (MERS-CoV) emerged in Saudi Arabia with 858 associated deaths by the end of November 2019 [3]. On December 12, 2019, first patient of COVID-19 was reported infected with SARS-nCoV2 strain in Wuhan-Hubei, China [4]. This infection got widespread and of May 01, 2020, 2,036,770 active cases, 234,279 (7.06%) deceased and 1,048,807 (31.59%) recovered has been reported worldwide [5]. This strain mainly belongs to the B-beta coronaviruses genus [6,7]. It has been observed that mortality rate vary from country to country which pondered the scientists to look into linkage between different variants of the SARS-nCoV2 with its severity along with other influencing factors e.g. temperature, testing facility, lockdown measures, aging factor and hygienic practices. Recently published genomic characterization study speculated the proximal origin of human Corona virus from Bat (*Rhinolophus affinis)* and Pangolin (*Maris javinica*) strains by natural selection in an animal host before zoonotic transfer or natural selection in humans following zoonotic transfer due to the novel observed variants in Receptor Binding Domain (RBD) and polybasic cleavage site of the Spike region [8].

In this comparative genomics study, phylogenetic analysis was carried out with 784 whole genome sequences available from NCBI Virus database in order to trace the closest Pakistani Corona virus homologues and then subject those closest strains for downstream processing. In an earlier analysis two Pakistani strains Gilgit1 MT240479 and Manga MT262993 were found in close proximity to USA MT263429 and China MT259229 strains. Further variant calling analysis using Galaxy platform on these 4 strains was performed to predict the effect of novel variants of Pakistani strains on the severity of COVID-19. SWISS-MODEL 3D structural modelling analysis was also performed for comparing mutant proteins of Pakistani strains to have an insight of potential role of these mutant proteins on pathogenicity of SARS-nCoV2.

## Methods

### Phylogenetic analysis

Rectangular cladogram was constructed using online NCBI Virus database phylogenetic tool [9] based on genbank sequence type and whole genome sequenced data taxid: 2697049 of 784 SARS-nCoV2 strains from all around the globe.

### Whole genome variant calling analysis

FASTA sequences of two Pakistan (Gilgit1 MT240479 and Manga MT262993), China (Wuhan) MT259229 and USA (Washington) MT263429 strains were retrieved from NCBI Genbank [10]. Pakistani FASTA sequences were converted to FastQ format using FASTA-to-Tabular-to-FASTQ tools (Galaxy Version 1.1.0) [11] and then mapped against reference genome using BWA MEM v 0.7.17.1 [12]. Mapped reads were coordinate sorted using SortSam feature and duplicate sequences were marked using MarkDuplicate feature of Picard tool. Aligned sequencing reads were processed for per position variant call using Naive Variant Caller (NVC) v 0.0.3 [13]. SnpSift Variant type [14] and SnpEff eff was used to annotate variants by custom building of reference sequence databases using SnpEff build v 4.3+T.galaxy4 (Figure. 1) [15].

**Figure 1.**
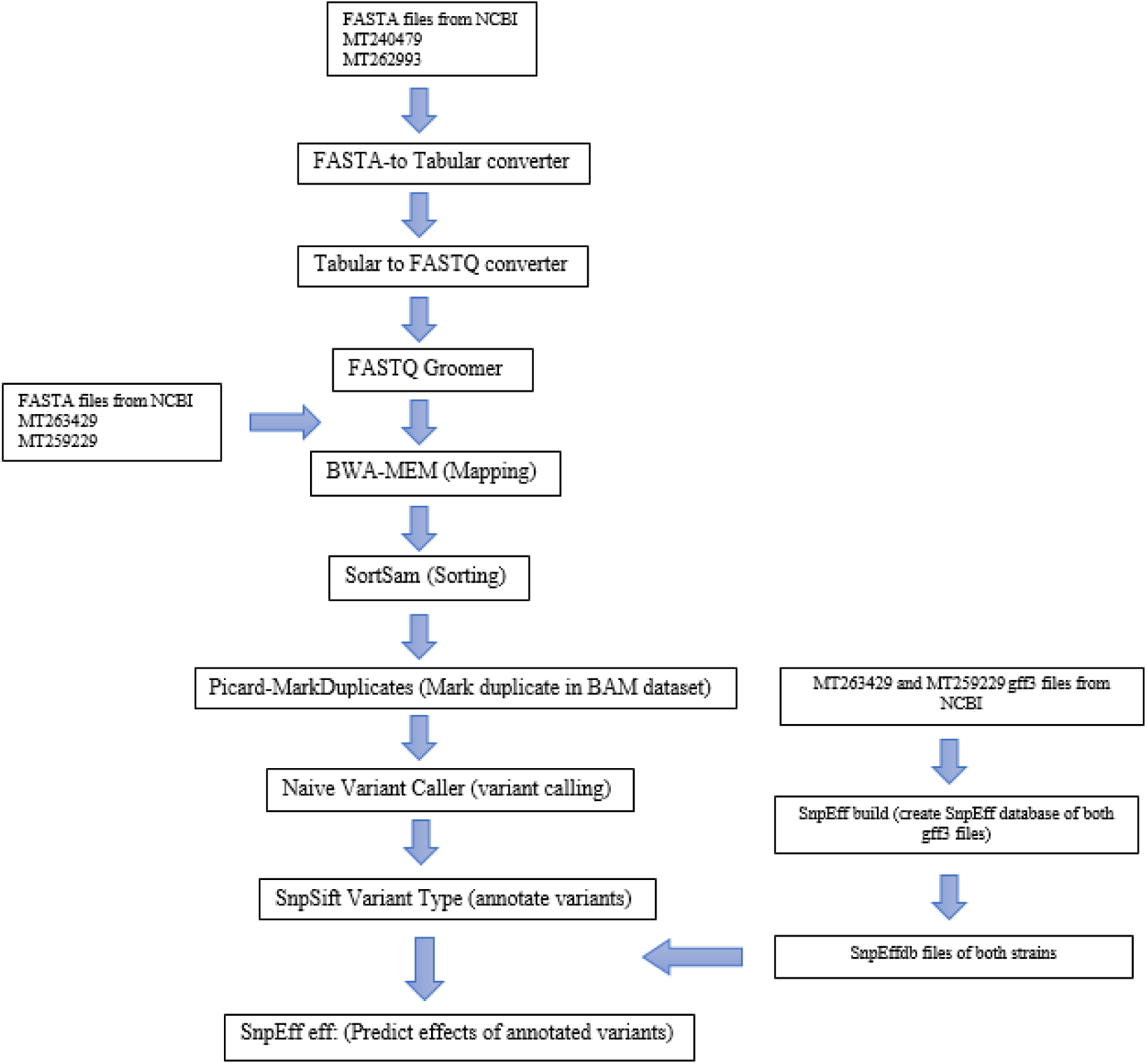
Variant calling workflow.

### Protein structure modelling

Proteins bearing variation in Pakistani strains were subjected to homology based structure modelling using Promod3 on Swiss model platform [16,17]. MT259229 (Chinese) and MT263429 (USA) were first subjected to search for homologues templates from which a prophesied 3D model was built. Next PDB files of these were used as template to build Pakistani mutant protein (N and ORF1a) 3D models based on target-template alignment along with Quaternary Structure Quality Estimate (QSQE) score complementing the GMQE for tertiary structure evaluation which is accomplished by supervised built-in “Support Vector Machine” (SVM) algorithm [18,19].

## Results

### Phylogenetic analysis

Cladogram constructed with NCBI Virus database showed relatedness of Pakistani MT240479 (Gilgit1) with MT184913 (CruiseA-USA) that shared same internal node from which the distance of MT240279 is 0.002342 and of MT184913 is 0.002381, while MT262993 (Manga-Pak) was present adjacent to MT039887 (WI-USA). MT262993 distance from its internal node is 0.000919 and that of MT039887 is 0.000865 (Figure. 2). Viruses in cladogram were from six continents including sequences from Africa (South Africa=1), North America (USA=680), Europe (Italy=2, Spain=11, France=1, Greece=4, Sweden=1), Asia (Pakistan=2, China=62, Iran=1, Turkey=1, India=2, Israel=2, Vietnam=2, Nepal=1, Taiwan=3, South Korea=4), Oceania (Australia=1) and from South America (1 strain each from Columbia, Peru and Brazil).

**Figure 2.**
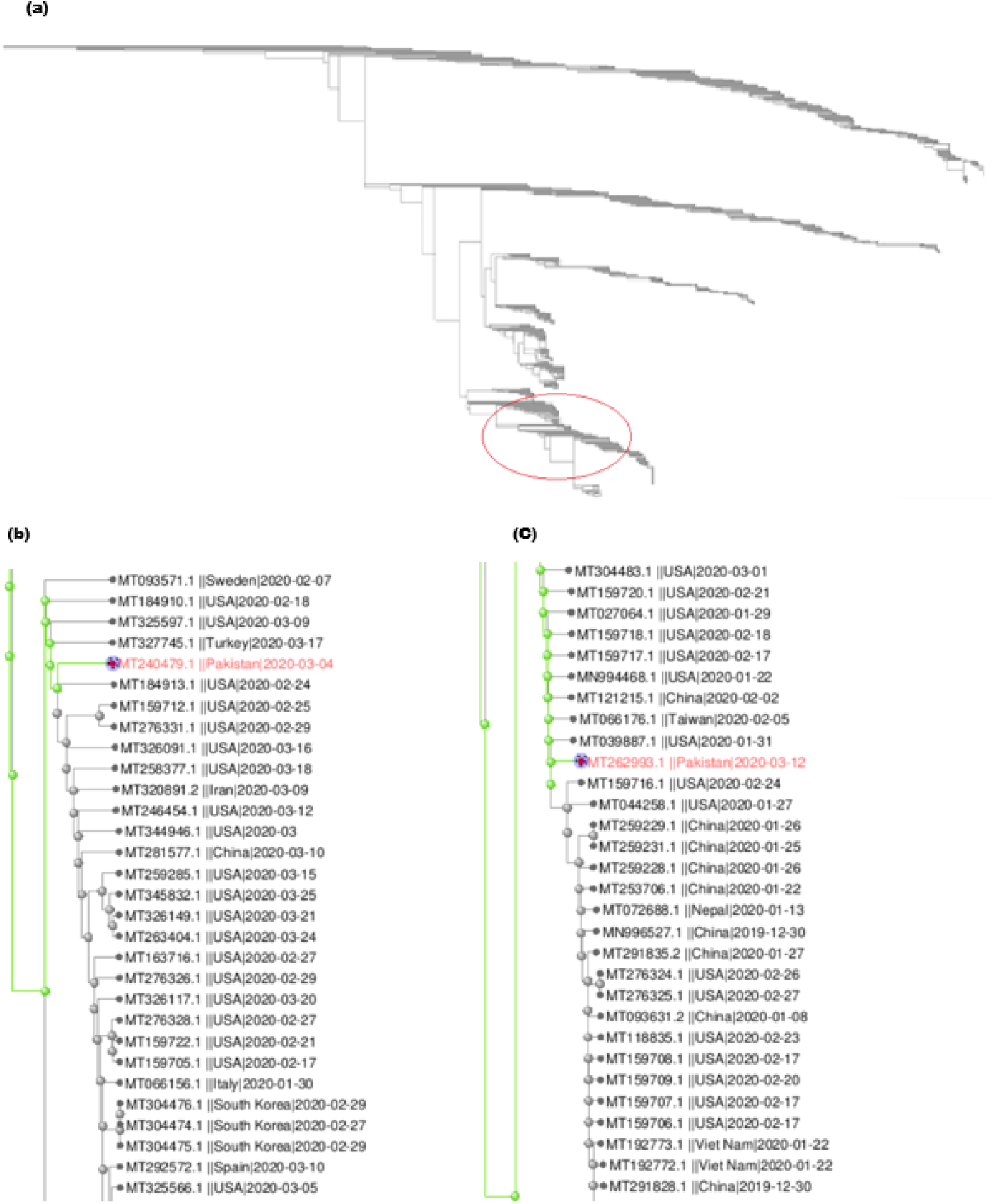
Overview of phylogenetic tree of 784 SARS-nCoV2 whole genome sequences. Red highlighted circle showing the clade of Pakistani strains (a), where red MT240479 (Gilgit1-Pak) strain is zoomed-in (b), and red MT62993 (Manga-Pak) strain is zoomed-in (c), presenting relatedness with other strains.

### Whole genome variant calling analysis of SARS-nCoV2

Pakistani strain MT262993 (29,836 bp) is more closely related to the virus strain MT259229 (28,964 bp) from China with the difference of 4 variations and each variant occurs on an average distance of 7,466 bp (Table 1), while MT263429 (29,889 bp) strain from USA is different with 9 variants loci at the rate of 1 variant per 2,988 bp (Table 2).

**Table 1.**
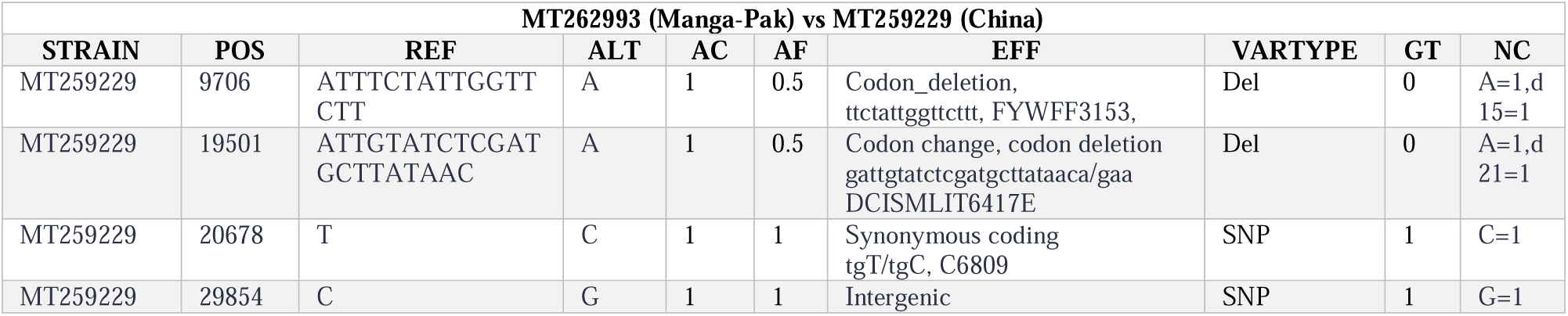
Detail of different variants observed in Manga-Pak strain and Chinese strain.

**Table 2.**
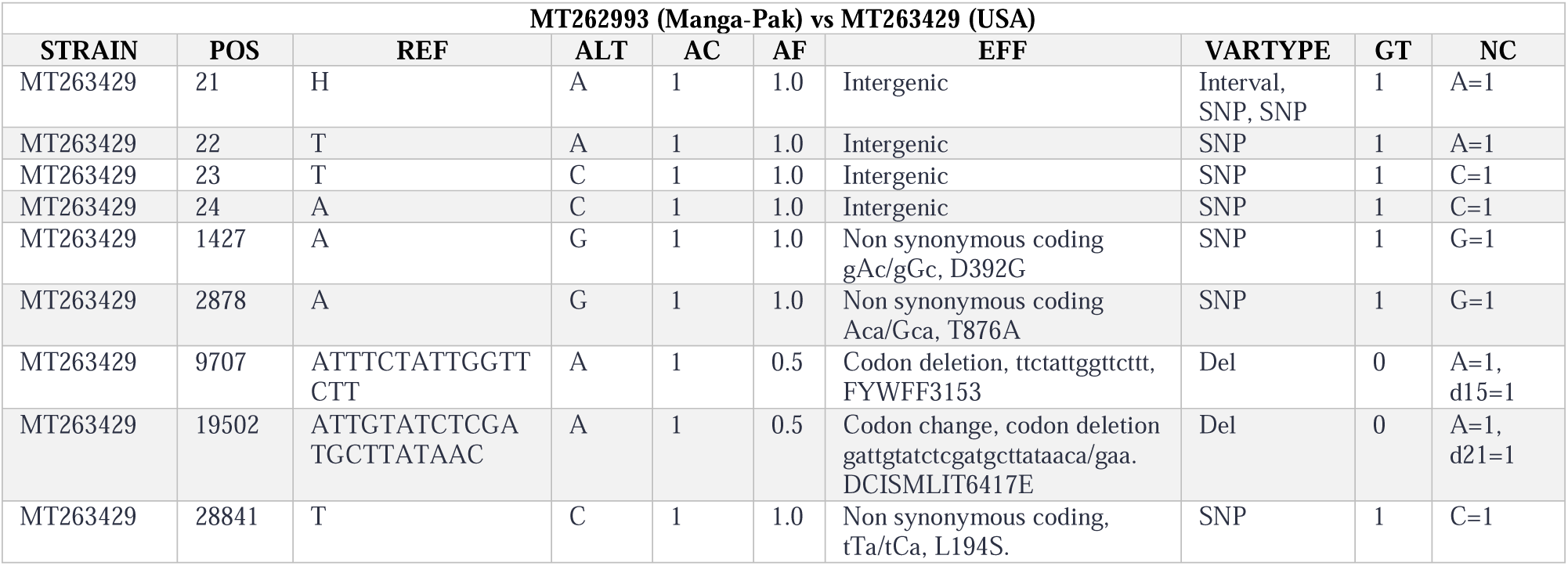
Detail of different variants observed in Manga-Pak and USA strain.

Gilgit1 MT240479 strain differ from MT259229 (Chinese) strain at 8 loci (SNPs) (Table 3), variants occurring after 3,733 bp while MT263429 (USA) differ from 10 variants (SNPs) each after 2,988 bp (Table 4).

**Table 3.** Detail of different variants observed in Gilgit1-Pak and Chinese strain.

| MT240479 (Gilgit1-Pak) vs MT259229 (China) |  |  |  |  |  |  |  |  |  |
| --- | --- | --- | --- | --- | --- | --- | --- | --- | --- |
| STRAIN | POS | REF | ALT | AC | AF | EFF | VARTYPE | GT | NC |
| MT259229 | 227 | C | T | 1 | 1 | Intergenic | SNP | 1 | T=1 |
| MT259229 | 870 | C | T | 1 | 1 | Non synonymous coding, Cgt/Tgt, R207C | SNP | 1 | T=1 |
| MT259229 | 1334 | C | T | 1 | 1 | Synonymous coding, ccC/ccT, P361 | SNP | 1 | T=1 |
| MT259229 | 1383 | G | A | 1 | 1 | Non synonymous coding, Gta/Ata, V378I | SNP | 1 | A=1 |
| MT259229 | 9145 | C | T | 1 | 1 | Non synonymous coding, cCt/cTt, P2965L | SNP | 1 | T=1 |
| MT259229 | 11069 | G | T | 1 | 1 | Non synonymous coding, ttG/ttT, L3606F | SNP | 1 | T=1 |
| MT259229 | 20678 | T | C | 1 | 1 | Synonymous coding, tgT/tgC/C6809 | SNP | 1 | C=1 |
| MT259229 | 29854 | C | G | 1 | 1 | Intergenic | SNP | 1 | G=1 |

**Table 4.** Detail of different variants observed in Gilgit1-Pak and USA strain.

| MT240479 (Gilgit1-Pak) vs MT263429 (USA) |  |  |  |  |  |  |  |  |  |
| --- | --- | --- | --- | --- | --- | --- | --- | --- | --- |
| STRAIN | POS | REF | ALT | AC | AF | EFF | VARTYPE | GT | NC |
| MT263429 | 24 | A | C | 1 | 1.0 | Intergenic | SNP | 1 | C=1 |
| MT263429 | 228 | C | T | 1 | 1.0 | Intergenic | SNP | 1 | T=1 |
| MT263429 | 871 | C | T | 1 | 1.0 | Non synonymous coding, Cgt/Tgt, R207C | SNP | 1 | T=1 |
| MT263429 | 1335 | C | T | 1 | 1.0 | Synonymous_coding, ccC/ccT, P361 | SNP | 1 | T=1 |
| MT263429 | 1384 | G | A | 1 | 1.0 | Non synonymous coding, Gta/Ata, V378I | SNP | 1 | A=1 |
| MT263429 | 1427 | A | G | 1 | 1.0 | Non synonymous coding, gAc/gGc, D392G | SNP | 1 | G=1 |
| MT263429 | 2878 | A | G | 1 | 1.0 | Non synonymous coding, Aca/Gca, T876A | SNP | 1 | G=1 |
| MT263429 | 9146 | C | T | 1 | 1.0 | Non synonymous coding, cCt/cTt, P2965L | SNP | 1 | T=1 |
| MT263429 | 11070 | G | T | 1 | 1.0 | Non synonymous coding, ttG/ttT, L3606F | SNP | 1 | T=1 |

The effects of variations by functional class for MT262993 (Manga) vs MT263429 (USA) are 05 missense variants, for MT240479 (Gilgit1) vs MT263429 (USA) are 13 missense and 02 silent while MT240479 vs MT259229 are 08 missense and 03 silent variations and 01 silent variant of MT262993 vs MT259229. Distribution of all variants impact, type and region are demonstrated in (Table 5). and the graphical representation of variants impacts is shown in (Figure. 3).

**Table 5.** Detail of Pakistani variants, their impact, type and region

| MT240479 (Gilgit1-Pak) vs MT263429 (USA) |  |  |  |  |  |
| --- | --- | --- | --- | --- | --- |
| Type |  | Count | Percent |  |  |
| Low |  | 2 | 11.765 |  |  |
| Moderate |  | 13 | 76.471 |  |  |
| Modifier |  | 2 | 11.765 |  |  |
| Type |  |  | Region |  |  |
| Type | Count | Percent | Type | Count | Percent |
| Intergenic | 2 | 11.765 | Exon | 15 | 88.235 |
| Non synonymous coding | 13 | 76.471 | Intergenic | 2 | 11.765 |
| Synonymous | 2 | 11.765 |  |  |  |
| MT262993 (Manga-Pak) vs MT263429 (USA) |  |  |  |  |  |
| Type |  | Count | Percent |  |  |
| Moderate |  | 8 | 61.538 |  |  |
| Modifier |  | 5 | 38.482 |  |  |
| Type |  |  | Region |  |  |
| Type | Count | Percent | Type | Count | Percent |
| Codon change, codon deletion | 1 | 7.602 | Exon | 8 | 61.538 |
| Codon deletion | 2 | 15.385 | Intergenic | 5 | 38.482 |
| Intergenic | 5 | 38.462 |  |  |  |
| Non synonymous coding | 5 | 38.462 |  |  |  |
| MT240479 (Gilgit1-Pak) vs MT259229 (China) |  |  |  |  |  |
| Type |  | Count | Percent |  |  |
| Low |  | 3 | 23.077 |  |  |
| Moderate |  | 8 | 61.538 |  |  |
| Modifier |  | 2 | 15.385 |  |  |
| Type |  |  | Region |  |  |
| Type | Count | Percent | Type | Count | Percent |
| Intergenic | 2 | 15.385 | Exon | 11 | 84.615 |
| Non synonymous coding | 8 | 61.538 | Intergenic | 2 | 15.385 |
| Synonymous | 3 | 23.077 |  |  |  |
| MT262993 (Manga-Pak) vs MT259229 (China) |  |  |  |  |  |
| Type |  | Count | Percent |  |  |
| Low |  | 1 | 20 |  |  |

| Moderate |  | 3 |  | 60 |  |
| --- | --- | --- | --- | --- | --- |
| Modifier |  | 1 |  | 20 |  |
| Type | Count | Percentage | Type | Count | Percent |
| Codon change, codon deletion | 1 | 20 | Exon | 4 | 80 |
| Codon deletion | 2 | 40 | Intergenic | 1 | 20 |
| Intergenic | 1 | 20 |  |  |  |
| Non synonymous coding | 1 | 20 |  |  |  |

**Figure 3.**
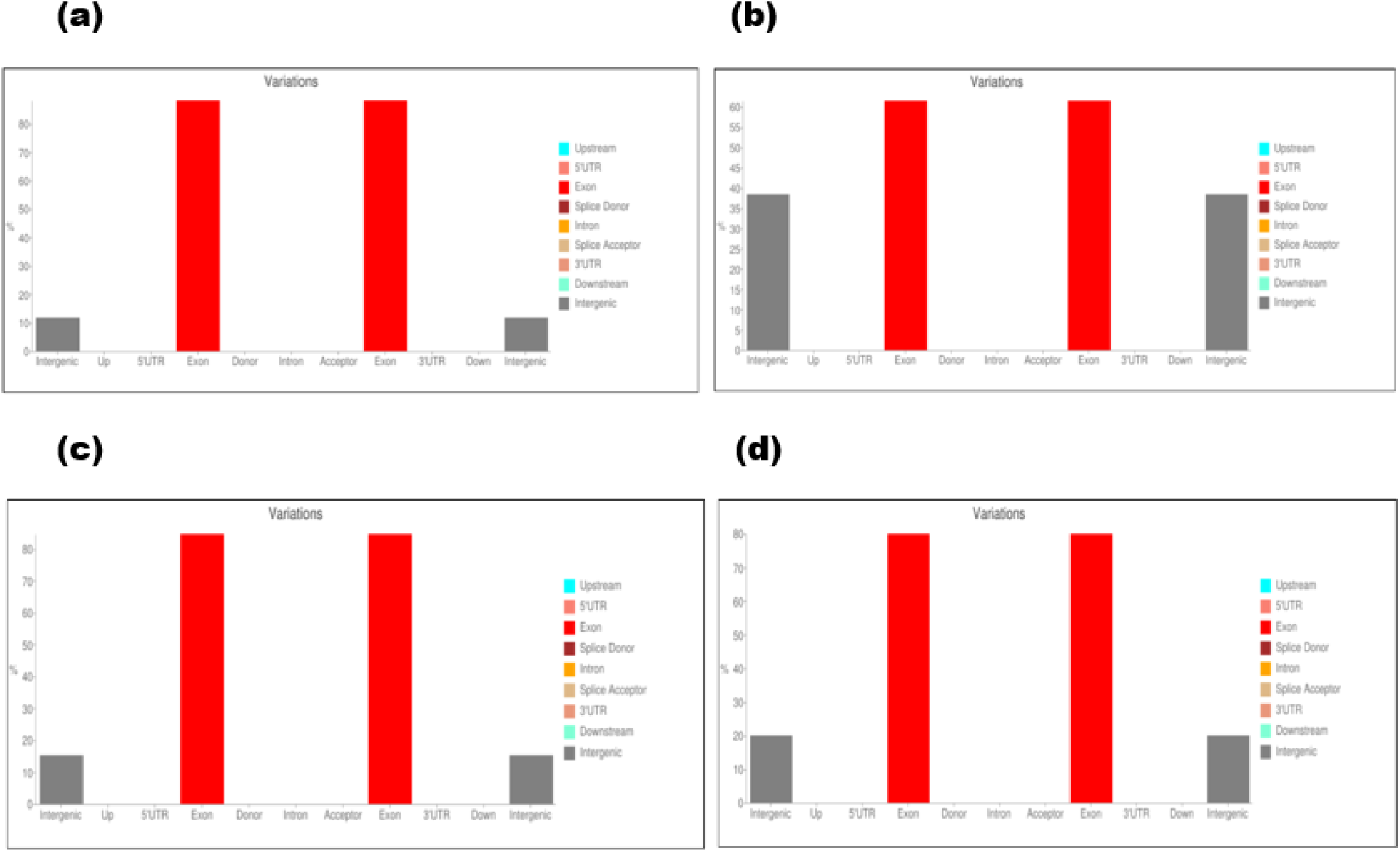
Graphic illustration of variatns type on x-axis and its occurance percentage on y-axis. MT240479 (Gilgit1) vs MT263429 (USA) (a), MT262993 (Manga) vs MT263429 (USA) (b), MT240479 (Gilgit1) vs MT259229 (China) (c), MT262993 (Manga) vs MT259229 (China) (d). Grey bar showing intergenic variants and red bar of exonic variants.

### Alteration in Pakistani SARS-nCoV2 genes and their effects

Detailed analysis of SARS-nCoV2 genome revealed some key alterations occurring in the sequences that codes for ORF1a polyprotein (4405aa) by cds-QIS61085.1 gene and by cds-QIS30017.1 gene. ORF1ab polyprotein (7096aa) by cds-QIS30016.1 gene and *ORF1ab* gene and Nucleocapsid phosphoprotein (419aa) expressed by *N* gene (Tables 6 and 7).

**Table 6.**
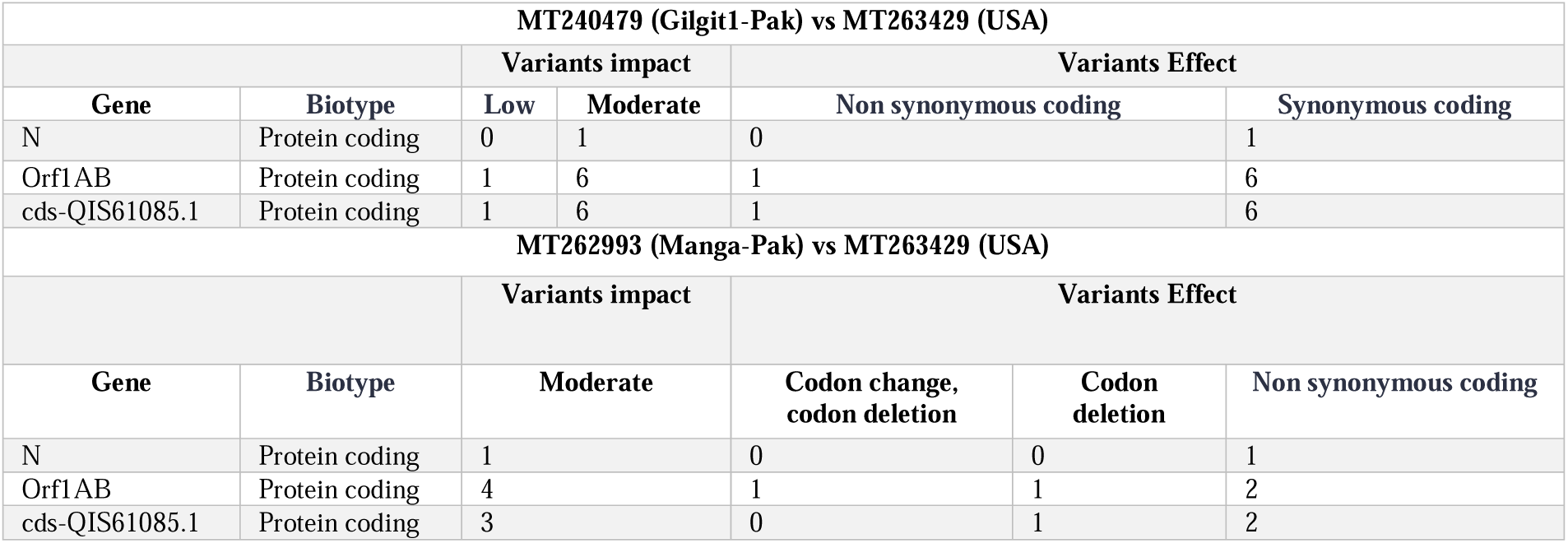
Altered genes of MT240479 (Gilgit1-Pak) and MT262993 (Manga-Pak) vs USA strain.

**Table 7.**
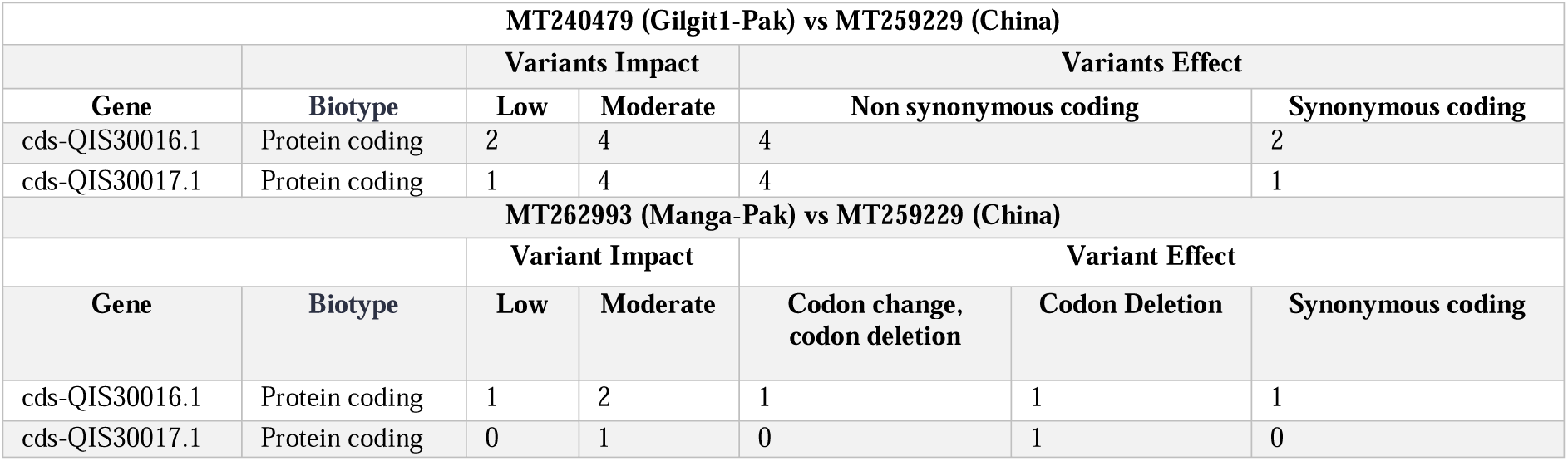
Altered genes of MT240479 (Gilgit1-Pak) and MT262993 (Manga-Pak) vs MT259229 (Chinese) strain.

N protein has multifarious activities, having 3 major domains of M-M (Matrix protein), M-N and N-S (S-spike protein) interaction [20]. It binds RNA tightly and packages the viral genome into capsid (a ribonucleoprotein) [21]. ORF1ab (Replicase polyprotein 1) encodes 7096 amino acids present at 5’ end and is involved in replication and translation of viral RNAs [22]. It interacts with the host innate immune response and is responsible for host virulence. Results also indicated that ORF1a polyprotein (4405aa) shows a rate of nonsynonymous substitutions usually [23].

### Pakistani CoV2 alteration on codon and amino acid level

Quantity and distribution of SNPs and deletions against USA and Chinese strains are shown in (Figure. 4). Genotypes of all samples were heterozygous. We considered relationship between codon usage, base substitutions SNVs and base deletions in annotated VCFs (Figure. 5).

**Figure 4.**
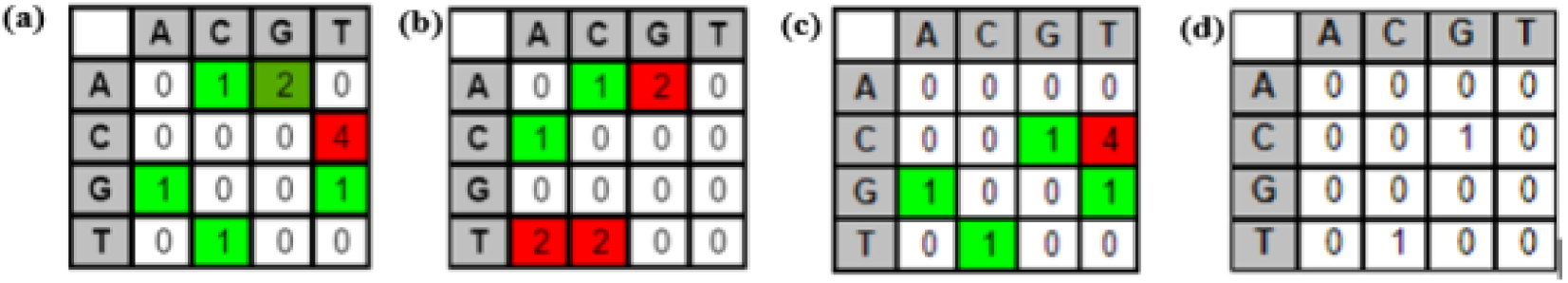
Comparision between MT240479 (Gilgit1-Pak) vs MT263429 (USA) having 10 SNPs with 8 transitions (Ts) and 2 transversions (Tv) (a), MT262993 (Manga-Pak) vs MT263429 (USA), 8 SNPs 4Ts, 2Tv + deletions (b), MT240479 (Gilgit1-Pak) vs MT259229 (China) 8 SNPs, 6Ts, 2Tv (c), and MT262993 (Manga-Pak) vs MT259229 (China) strain of SARS CoV-2 displaying 2 bp changes (SNPs), 1 Ts and 1 Tv (d).

**Figure 5.**
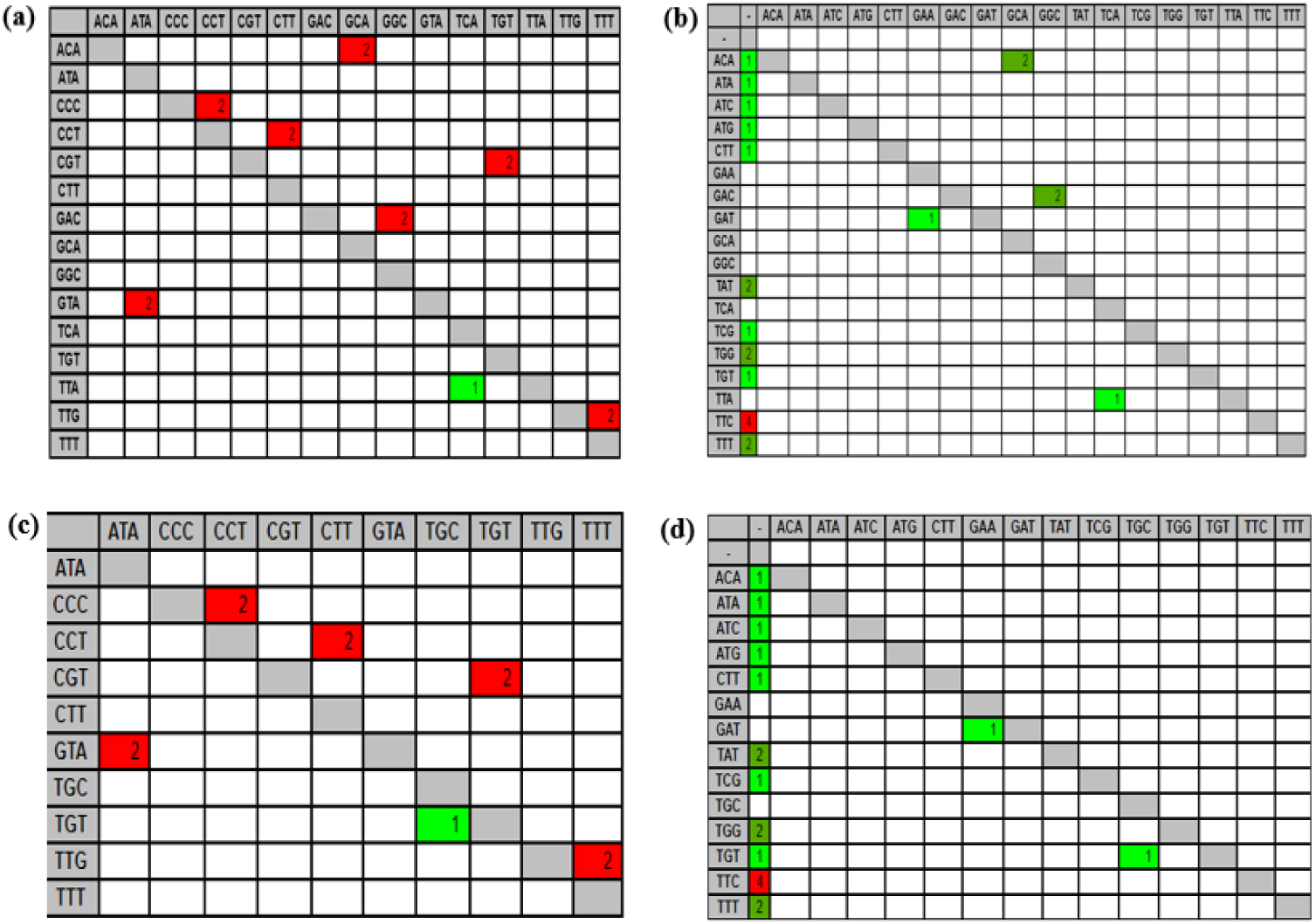
Results of codon changes. MT240479 (Gilgit1-Pak) vs MT263429 (USA) (a), MT262993 (Manga-Pak) vs MT263429 (USA) (b), MT240479 (Gilgit1-Pak) vs MT259229 (China) (c), MT262993 (Manga-Pak) vs MT259229 (China) (d), strain of SARS CoV-2. Columns indicating changed codons as a result of variation. Red color is signifying heat-map i.e. higher variation.

Amino acid changes in annotated VCF were also identified. Surprisingly, there is a multiple nucleotide deletion in MT262993 strain TTTCTATTGGTTCTT (FYWFF3153), ATTGTATCTCGATGCTTATAAC (DCISMLIT6417E) at loci 9706, 19501 with MT259229 and at positions 9707, 19502 when comparing it with MT263429 that might be causing alteration in protein effectiveness (Figure. 6).

**Figure 6.**
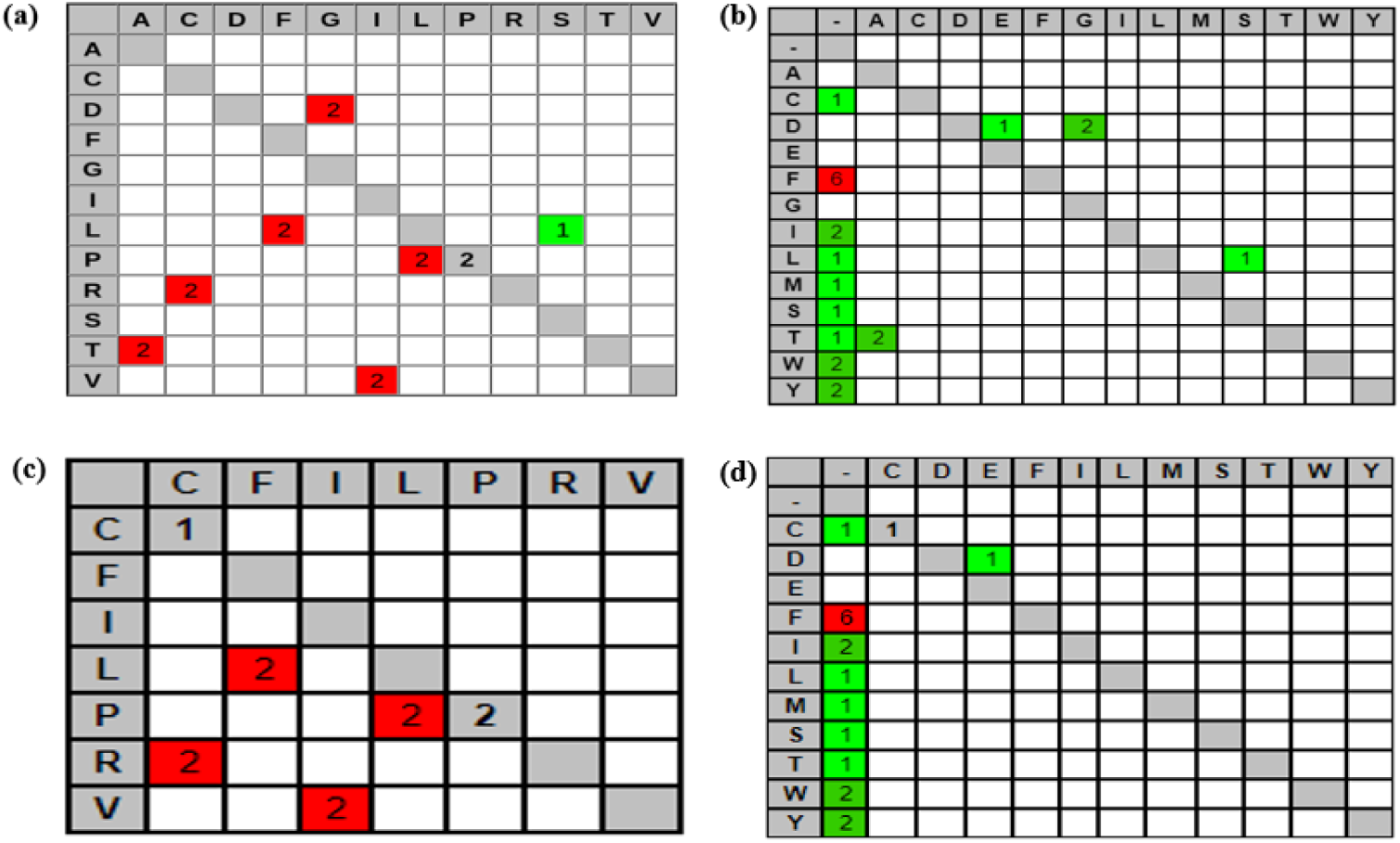
Detailed explanation of amino acid change in MT240479 (Gilgit1-Pak) vs MT263429 (USA) (a), MT262993 (Manga-Pak) vs MT263429 (USA) (b), MT240479 (Gilgit1-Pak) vs MT259229 (China) (c), MT262993 (Manga-Pak) vs MT259229 (China) (d), strains of SARS CoV-2. Where rows are reference codons and columns are changed amino acid. The count of impact is showing high variation. Red background color indicates that more changes happened.

### Variants distribution across the whole genome of Pakistani Corona virus strains

Division of variants across the whole genome of SARS-nCoV2 before 2900 bases are shown in (Figure. 7).

**Figure 7.**
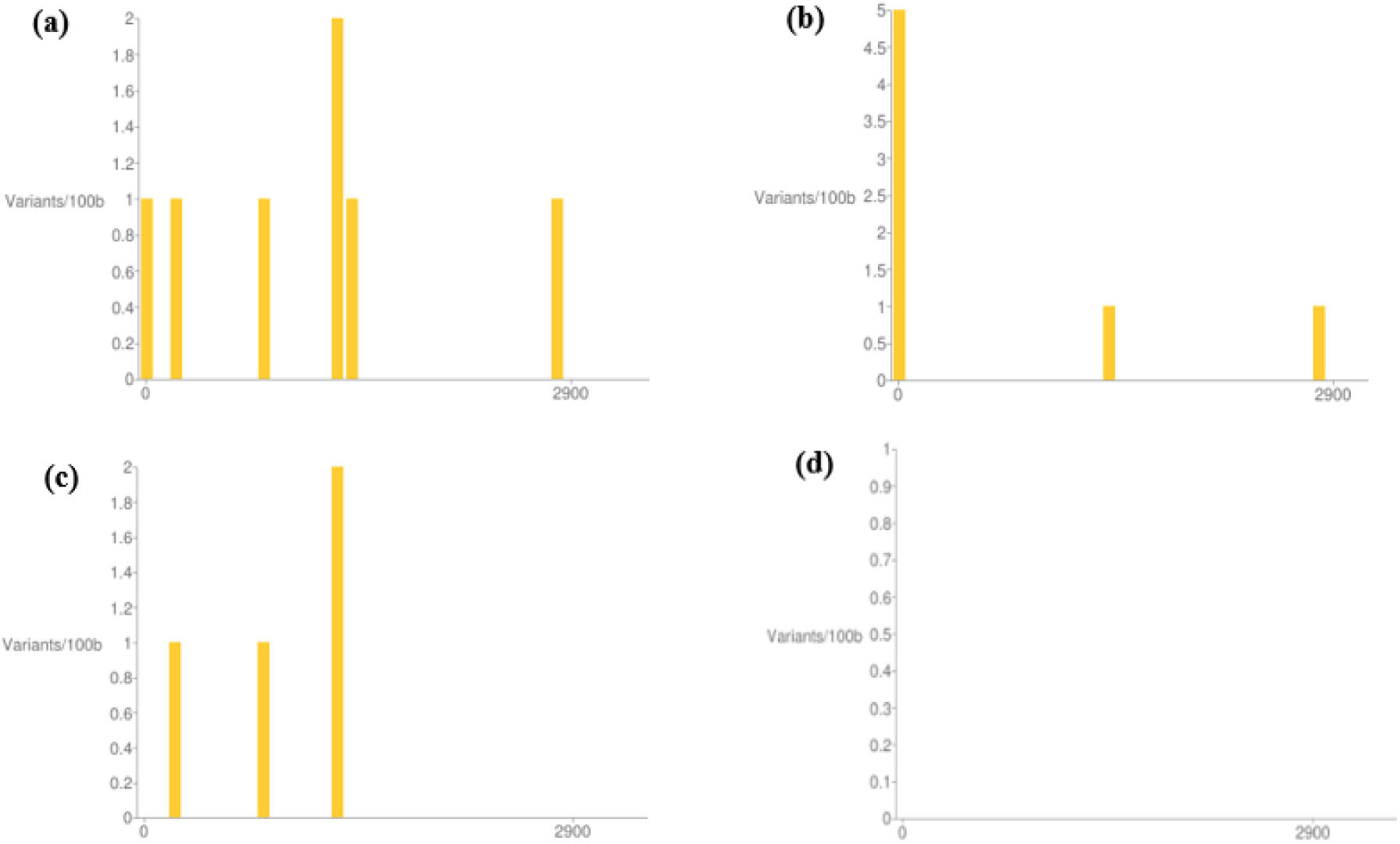
Bar chart illustrating variants across the whole genome (first 2900 bases are shown here). MT240479 (Gilgit1-Pak) vs MT263429 (USA) (a), MT262993 (Manga-Pak) vs MT263429 (USA) (b), MT240479 (Gilgit1-Pak) vs MT259229 (China) (c), MT262993 (Manga-Pak) vs MT259229 (China) (d). x-axis showing bases and y-axis showing variants/100 bases.

Seven SNPs were observed in first 2900 bases while other 02 are present in the remaining genome between Pakistani Manga and USA strain, while Pakistani Gilgit1 strain has total 10 SNPs, first seven within 2900 bases are shown in (Figure. 7b). Total eight variants appeared in Gilgit1 vs Chinese strain, first four variants are shown in (Figure. 7c). Similarly total 04 variants (SNPs=2, del=2), none of the polymorphism within first 2900 bases are observed in Manga vs Chinese strain shown in (Figure. 7d), exact positions and genes are mentioned in (Tables 1-4, 6, 7)

### Homology modelling of Pakistani SARS-nCoV2 mutant proteins

N protein which has mutations when Pakistani strains were compared to MT263429 USA was modelled from partial alignment and structurally analyzed. Comparison between Gilgit1 and USA has sequence similarity of 62% with 26% sequence coverage, aligning amino acids in the range of 250-360 from total 419aa, while Manga strain has sequence similarity with USA of 63%, sequence coverage 30% aligning 46-172 amino acids out of 419aa (Figure. 8).

**Figure 8.**
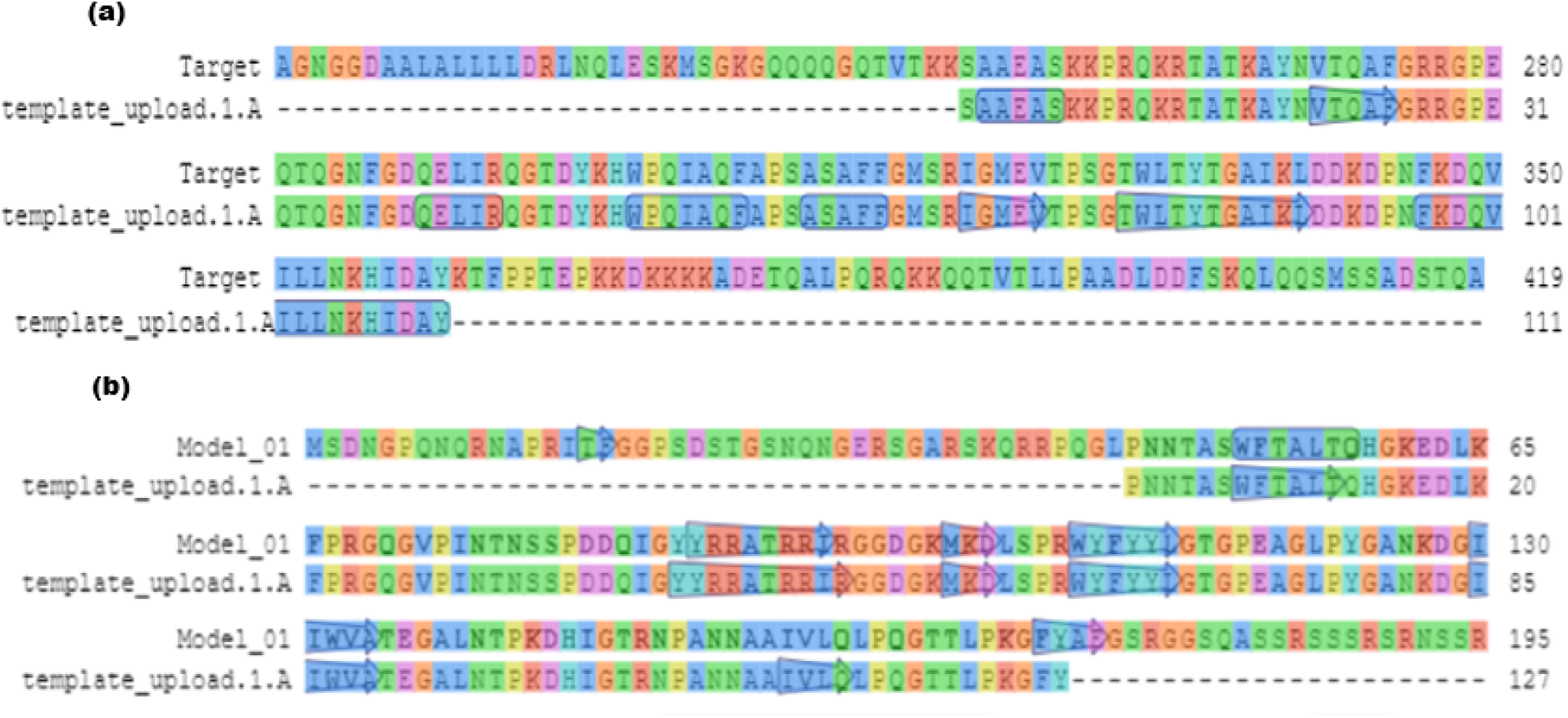
Target-template alignment of N protein. 250-360 amino acids aligned between target MT240479 (Gilgit1-Pak) and template MT263429 (USA) (a), 46-172 amino acids are aligned between target MT262993 (Manga-Pak) and template MT263429 (USA) (b).

MT240479 Gilgit1 has more loop structures than MT262993 Manga N protein and no ligand interaction domains were found in template (USA) (Figure. 9).

**Figure 9.**
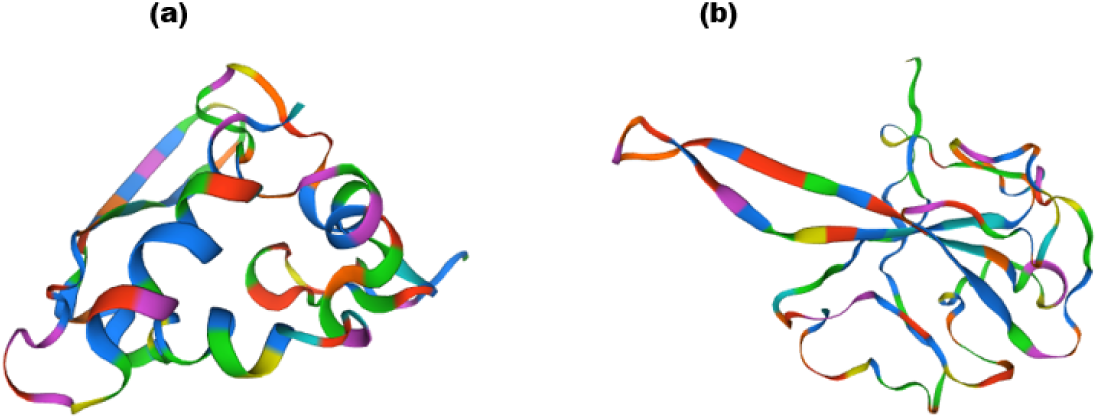
Protein 3D structures modelled. MT240479 (Gilgit1-Pak) N protein (a), and MT262993 (Manga-Pak) N protein (b). For both MT263429 (USA) was used as template.

ORF1ab protein amino acid sequence was too large to be modelled. At last, mutant ORF1a proteins were then structurally modelled by using both Chinese and USA strain sequences as template. Partial alignment of 1567-1878 amino acids (7% of total sequence coverage) among Gilgit1 and Manga strains were compared against USA and Chinese strain resulted in 62% sequence similarity in all four alignments (Figure. 10).

**Figure 10.**
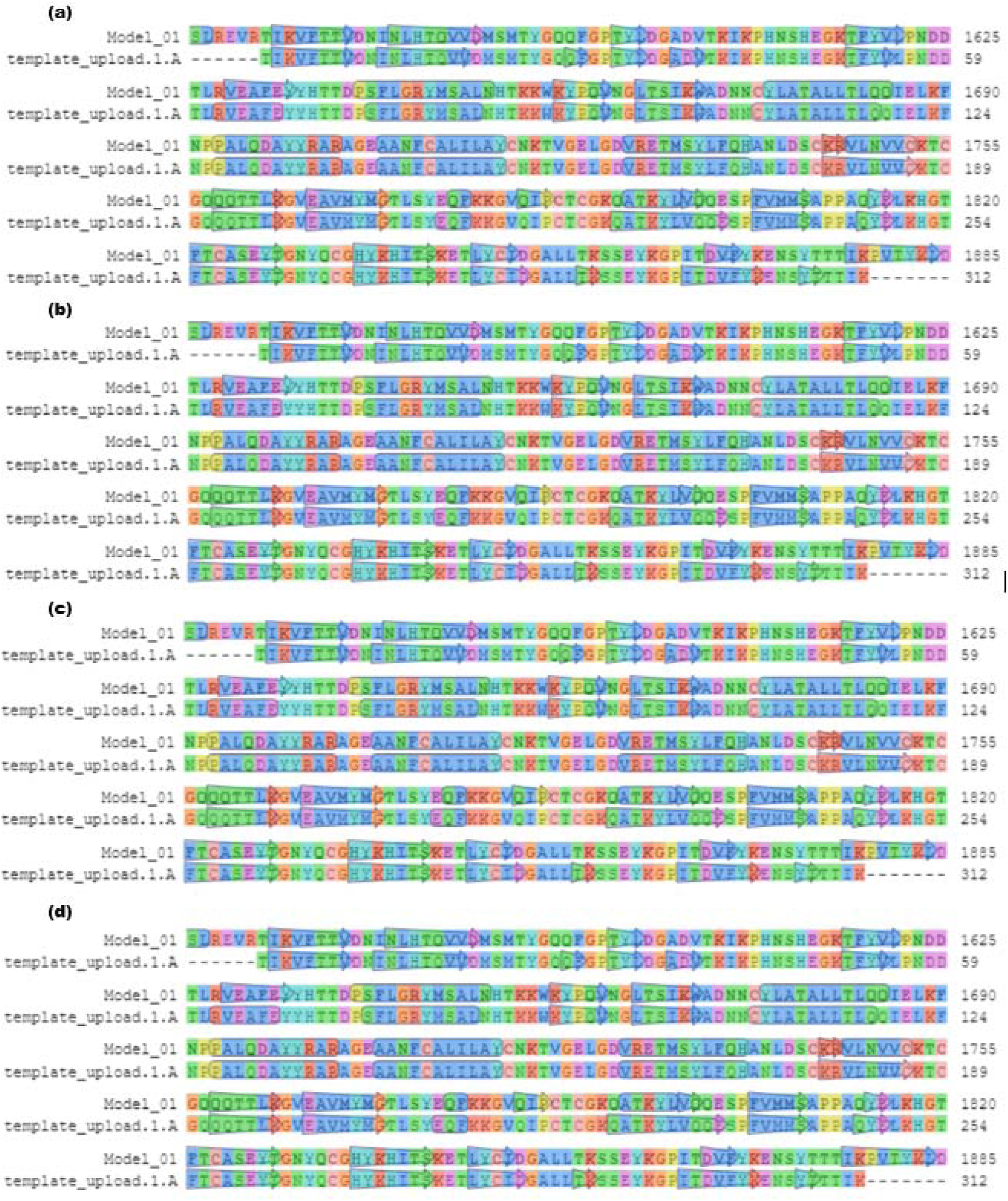
Target-template partial alignment. MT240479 (Gilgit1-Pak) vs MT263429 (USA) (a), MT262993 (Manga-Pak) vs MT263429 (USA) (b), MT240479 (Gilgit1-Pak) vs MT259229 (China) (c), MT262993 (Manga-Pak) vs MT259229 (China) (d). Alignment from 1567-1878 amino acids out of 4405aa is shown in all four alignments.

Structures resulting from ORF1a protein alignment revealed three major non-covalent protein-ligand interaction domains in all ORF1a protein structures. Ligand Zn^+2^ forms interaction with their chain A (Figure. 11).

**Figure 11.**
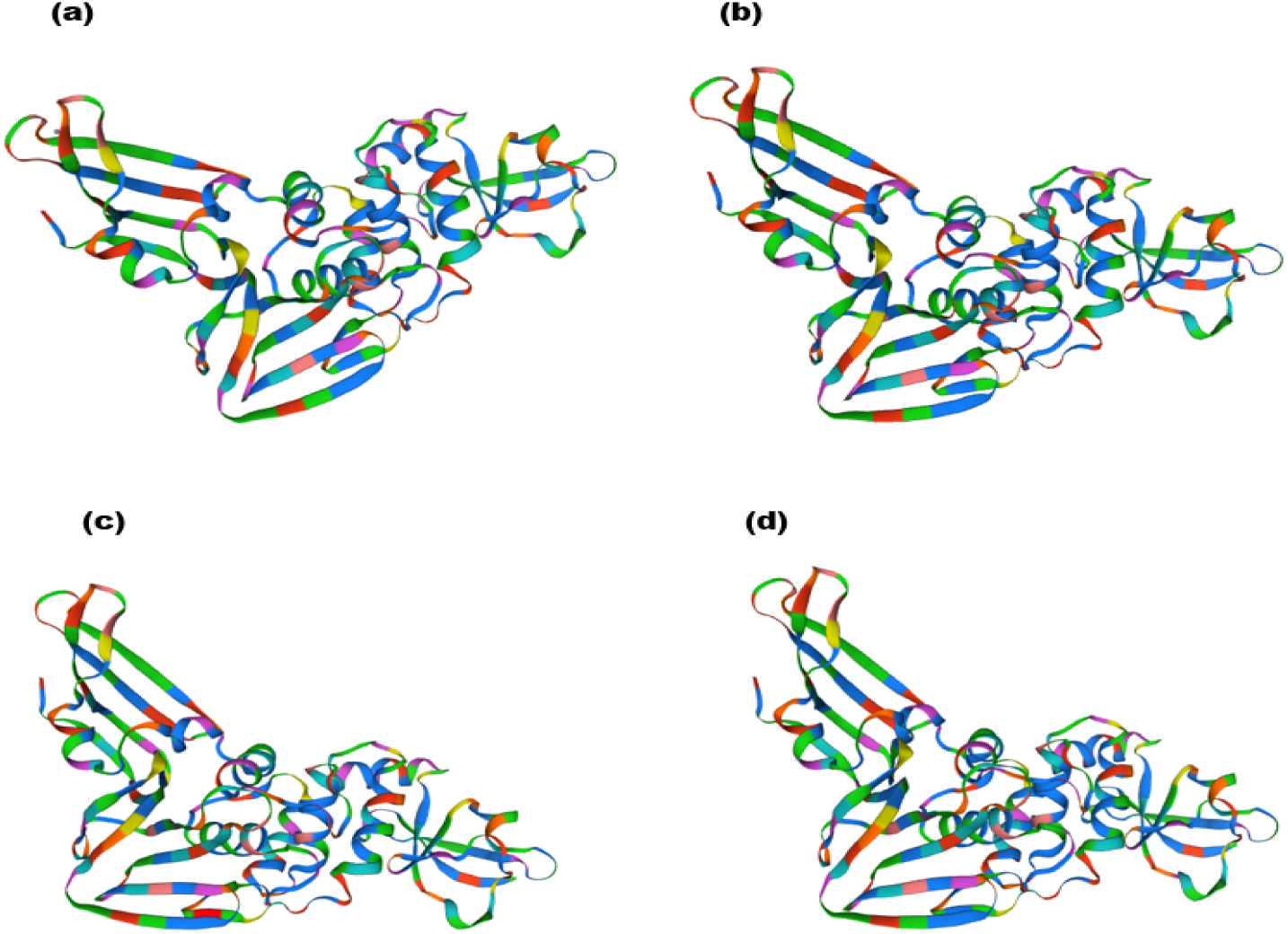
ORF1a protein structurally modelled MT240479 (Gilgit1-Pak) (a), and MT262993 (Manga-Pak) (b), both modelled using MT263429 (USA) as template. MT240479 (Gilgit1-Pak) (c), while MT262993 (Manga-Pak) (d), which were modelled against MT240479 (China) ORF1a protein.

## Discussion

Around 70% of human pathogens are of zoonotic origin including all Corona viruses types and one of them the SARS-nCoV2 also, which is causing COVID-19, the ongoing pandemic [24]. By today May 1, 2020 210 countries and territories are affected. In the current era Next Generation Sequencing (NGS) techniques are booming to let genomic data available seamlessly to scientists and researchers for deciphering an unseen in depth view of viruses on genomic level to overcome and combat this disease. For genomic level characterization and comparison, we did phylogenetic study on 784 Corona virus strains based on whole genome nucleotide sequenced available data, the two Pakistani strain MT240479 occurred in the same clade with MT184913 (CruiseA-USA) and MT262993 appeared in close proximity to MT039887 (WI-USA) strain. We also performed variant calling and detected 31 variants in *N, ORF1ab* and *ORF1a* genes while comparing Pakistani strains with MT263429 and MT259229. These mutational events come up with some predictive evidence about the outcomes of cases. On January 19, 2020 first case of COVID-19 was reported in Washington, USA and after one and half month time death rate was 27% while in Pakistan first case appeared on February 26, 2020 and death rate after the same time was 10.14% with 89.86% recovery rate, the death rate still going low up to today’s statistics. Similarly, China too suffered from very high ratio of deceased people 43.18% same time after its first case was being reported [25] Gilgit1 and Manga strain variants are assumed to be linked with low severity of this pandemic along with other factors of high temperature, less elderly people and ablution practices in Pakistan. Possibly variations in proteins of Pakistani SARS-nCoV2 might also have affected its interaction with the Angiotensin-converting enzyme 2 (ACE2) receptors in humans causing low virulence but for validation of this assumption functional studies are needed. We also performed structural analysis and modelled mutant proteins 3D structure from target-template alignment using N and ORF1a proteins of USA and Chinese strains as template and both Pakistani N and ORF1a as target sequences to visualize it on amino acid level to further ensure the differences in studied strains.

We conclude our discussion by making the instance that *N* and *ORF1a* genes variants in Pakistani Corona virus might be associated with some functional phenotype causing low mortality rate in Pakistan vs USA and Chinese strains, however no variants were found in RBD and polybasic cleavage site of spike region which is more critical region for the virulency of this virus. This hypothesis still needs more research to validate and to find out association of these mutant genes and other influencing factors with the pathogenicity of this virus.

## Acknowledgements

Authors are thankful to the University Institute of Biochemistry and Biotechnology, Pir Mehr Ali Shah Arid Agriculture University, Rawalpindi and Department of Healthcare Biotechnology, National University of Sciences and Technology (NUST), Islamabad, Pakistan for their generous sequencing efforts and making their data publically available on NCBI to facilitate the researcher community.

